# Metabolic flux regulates growth transitions and antibiotic tolerance in uropathogenic *Escherichia coli*

**DOI:** 10.1101/2023.05.09.540013

**Authors:** Josiah J. Morrison, Daniel A. Banas, Ellen K. Madden, Eric C. DiBiasio, David C. Rowley, Paul S Cohen, Jodi L. Camberg

## Abstract

Reducing growth and limiting metabolism are strategies that allow bacteria to survive exposure to environmental stress and antibiotics. During infection, uropathogenic *Escherichia coli* (UPEC) may enter a quiescent state that enables them to reemerge after completion of successful antibiotic treatment. Many clinical isolates, including the well characterized UPEC strain CFT073, also enter a metabolite-dependent, quiescent state in vitro that is reversible with cues, including peptidoglycan-derived peptides and amino acids. Here, we show that quiescent UPEC is antibiotic tolerant and demonstrate that metabolic flux in the tricarboxylic acid (TCA) cycle regulates the UPEC quiescent state via succinyl-CoA. We also demonstrate that the transcriptional regulator complex IHF and the FtsZ-interacting protein ZapE, which is important for *E. coli* division during stress, are essential for UPEC to enter the quiescent state. Notably, in addition to engaging FtsZ and late-stage cell division proteins, ZapE also interacts directly with TCA cycle enzymes in bacterial two hybrid assays. We report direct interactions between succinate dehydrogenase complex subunit SdhC, the late-stage cell division protein FtsN, and ZapE. These interactions likely enable communication between oxidative metabolism and the cell division machinery in UPEC. Moreover, these interactions are conserved in an *E. coli* K-12 strain. This work suggests that there is coordination among the two fundamental and essential pathways that regulate overall growth, quiescence, and antibiotic susceptibility.

**Importance:** Uropathogenic *Escherichia coli* (UPEC) are the leading cause of urinary tract infections (UTIs). Upon invasion into bladder epithelial cells, UPEC establish quiescent intracellular reservoirs that may lead to antibiotic tolerance and recurrent UTIs. Here, we demonstrate using an in vitro system that quiescent UPEC cells are tolerant to ampicillin and have decreased metabolism characterized by succinyl-CoA limitation. We identify the global regulator the IHF complex and the cell division protein ZapE as critical regulators of quiescence and antibiotic tolerance. Lastly, we show ZapE interacts with components of both the cell division machinery and the TCA cycle, and this interaction is conserved in non-pathogenic *E. coli*, establishing a novel link between cell division and metabolism.

## Introduction

Mechanisms used by bacteria to subvert the effects of antibiotic treatment include antibiotic resistance, which occurs through acquired genetic changes (1–3), persistence, which selects for low frequency bacterial survivors (4–7), and antibiotic tolerance, which can be evoked in cells through limited growth and metabolism (8–11). Reciprocally, activating cellular metabolism is known to increase susceptibility to antibiotics (12, 13). Bactericidal antibiotics including beta-lactams also appear to stimulate metabolic activity leading to production of toxic metabolites, thus enhancing their effect (14, 15). Consistent with the importance of metabolism in antibiotic-induced killing, a recent report showed that deletion of the *E. coli sucA* gene, which encodes a subunit of the tricarboxylic acid cycle (TCA) enzyme alpha ketoglutarate dehydrogenase, conferred high levels of protection against carbenicillin (16).

Uropathogenic *Escherichia coli* (UPEC) enter a dormant state in vitro and were suggested to populate quiescent intracellular reservoirs (QIRs) during infection in vivo. UPEC are the leading cause of urinary tract infections (UTIs) (17–19) and greater than 25% of women have a UTI that recurs within 6 months of the initial infection (20, 21). Of these, 77% are caused by the same strain of *E. coli* that initially caused the infection (22). QIRs are resistant to the host immune response and antibiotic treatment and thus may serve as a source of recurrent infection (23–25). Host cell differentiation and actin cytoskeletal arrangement may promote reinitiation of replication for these quiescent UPEC, and this may result in recurrence of UTIs (26, 27).

At low inoculation density (<10^6^ CFU ml^-1^), many clinical UPEC isolates, including the widely studied ST-73 strain CFT073, are non-proliferative, or quiescent, on minimal medium containing glucose as the sole carbon source (28, 29). At high inoculation density, i.e., > 10^6^ CFU ml^-1^, the quiescence-capable UPEC cells are proliferative on glucose minimal medium, suggesting that proliferation on glucose is quorum-dependent. Cues, including peptidoglycan tetra-and pentapeptides, and a combination of L-lysine and L-methionine override the requirement for a quorum and enable proliferation at low inoculation density, although the mechanism is unknown. A mini transposon screen uncovered several carbon metabolism genes, including *gnd*, *pykF*, *zwf*, *gdhA*, and *sdhA*, that are critical for establishing the quiescent state (29).

Here, we investigated the role of TCA cycle metabolism during quiescence in UPEC strain CFT073. We also discovered two additional genes, *ihfB* and *zapE*, which encode a transcriptional regulator and an FtsZ-binding protein, respectively, that are important for the quiescent state (30–32). We demonstrate that quiescent cells are antibiotic-tolerant in the presence of ampicillin, and this is modulated by succinate levels. Through genetic analysis, we demonstrate that the limitation of succinyl-CoA in cells is necessary to maintain the quiescent state, and this is likely regulated by the IHF transcriptional regulator. Lastly, we show that ZapE, which interacts with FtsZ and FtsN (33), also engages subunits of the succinate dehydrogenase complex in bacterial two hybrid assays. Together, our work demonstrates that the in vitro, quiescent state exhibited by the UPEC strain CFT073 confers antibiotic tolerance and is modulated by TCA cycle metabolism. As a key regulator of quiescence, ZapE communicates with cell division and metabolism complexes in the cell, and this may underlie how ZapE can prevent proliferation and promote quiescence. The protein-protein interactions among division and metabolism complexes reported here are preserved in *E. coli* K-12 strains, suggesting that coordination between metabolism and division is evolutionarily conserved.

## Results

### Mini-transposon screen reveals regulators of quiescence

To identify genes in UPEC that are essential for establishing the low-density quiescent state on glucose minimal agar, we performed a high throughput mini-transposon (Tn5) screen. We introduced a mini-Tn5 transposon in the UPEC genome (CFT073) by conjugative transfer from a donor strain (ATM161), and then screened a culture of recipients (10^4^ CFU ml^-1^) for growth on glucose minimal agar supplemented with kanamycin. We previously reported that UPEC cells (CFT073) become quiescent on glucose minimal agar at low plating density (<10^6^ CFU ml^-1^) but proliferate rapidly at high plating density (>10^6^ CFU ml^-1^) (28, 29). We isolated 21 mutant strains that formed colonies on glucose minimal agar when plated at low cells density (10^4^ CFU ml^-1^) and identified the mini-Tn5 insertion site by arbitrarily-primed PCR and sequencing (34). Of these mutant strains, 19 had transposon insertions in *gnd*, a gene that encodes 6-phosphogluconate dehydrogenase, an enzyme in the pentose phosphate pathway, and was previously implicated in quiescence (29) (Fig. S1A). In that study, *gnd,* along with four other genes involved in central carbon metabolism (*ghdA*, *sdhA*, *pykF*, and *zwf*) were identified. In the current study, we identified two additional strains defective for quiescence with mini-Tn5 insertions in genes that do not encode enzymes involved in carbon metabolism, *ihfB* and *zapE*. Both strains, (JM205) and (JM206), formed colonies on glucose minimal agar inoculated with 10^4^ CFU ml^-1^ (Fig. 1A) (Fig. S1A), in contrast to the wild type strain (CFT073), which is unable to proliferate at low cell density, but proliferates at high density (Fig. 1A) (Fig. S1A). The *ihfB* gene encodes the beta subunit of integration host factor (IHF). IHF is a DNA-binding transcriptional regulator complex comprising IhfB and IhfA (35). The IHF complex has been shown to be important for virulence of UPEC by pathogenicity island stabilization (36). The IHF regulon in *E. coli* (K-12) is predicted to contain more than 100 genes, including many involved in central carbon metabolism such as *gltA icd*, *aceA*, *sucA, sucB*, *sucC, and sucD* (37–39) (40, 41). The second gene identified as a site of mini-Tn5 insertion, *zapE*, encodes the cell division regulator ZapE, which binds to FtsZ polymers assembled with GTP and is important for cell growth during stress (31) (32).

**Figure 1.**
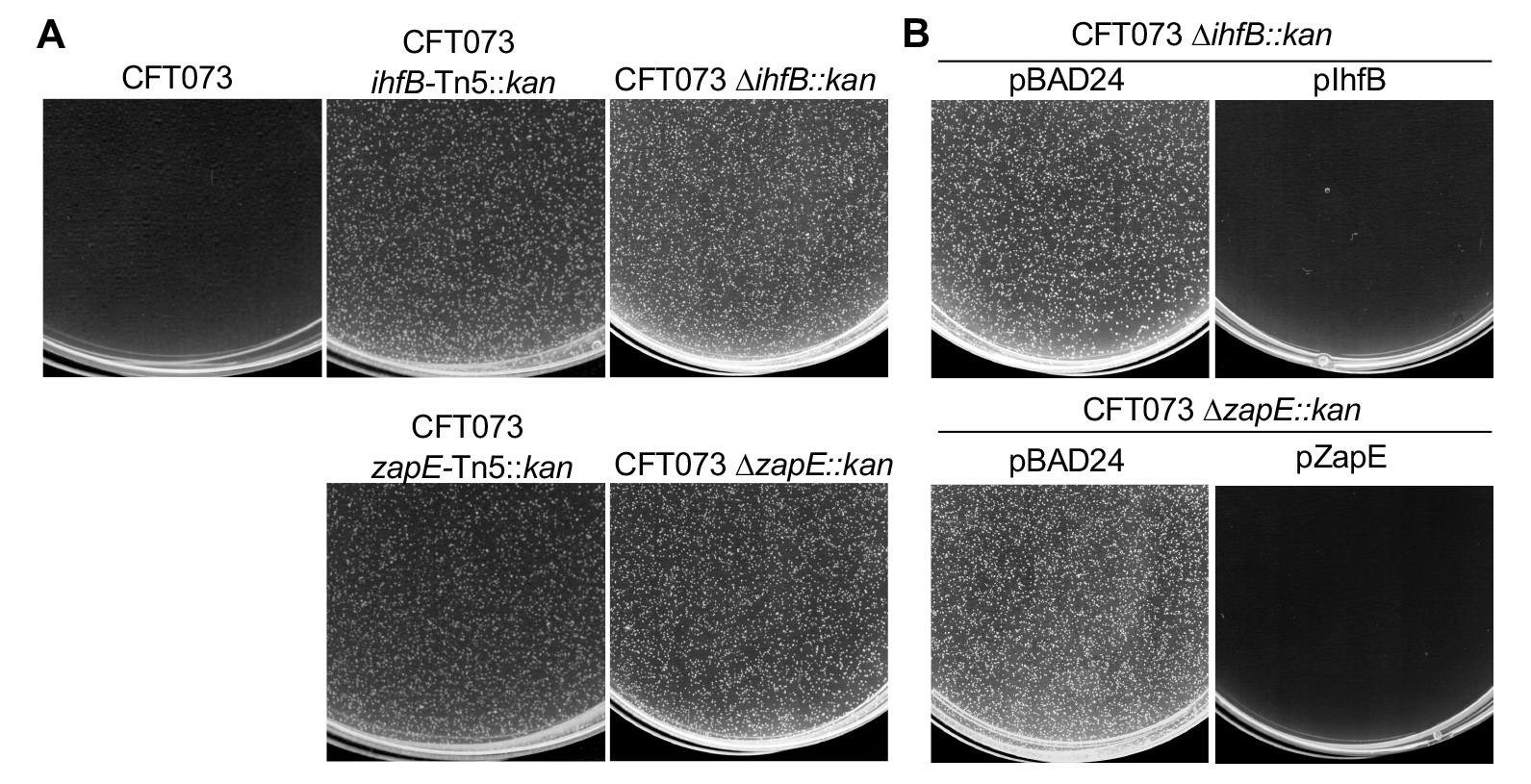
High-throughput screen for genes critical for UPEC CFT073 quiescence. (A) CFT073, CFT073 *ihfB* Tn5::*kan*, CFT073 *zapE* Tn5::*kan*, CFT073 Δ*zapE*, and CFT073 Δ*ihfB* were plated at 10^4^ CFU ml^-1^ on 0.2% glucose minimal agar and incubated at 37 °C for 24 h. (B) CFT073 Δ*ihfB* containing pBAD24 or pIhfB and CFT073 Δ*zapE* containing pBAD24 or pZapE were plated at 10^4^ CFU ml^-1^ on 0.2% glucose minimal agar and incubated at 37 °C for 24 h. CFT073 Δ*ihfB* and CFT073 Δ*zapE* containing pBAD24, pIhfB, or pZapE were supplemented with 0.1% arabinose and 50 μg ml^-1^ ampicillin. Pictures in (A) and (B) are representative of at least three independent experiments.

To confirm if *ihfB* and *zapE*, which were disrupted by the mini-Tn5, are essential for quiescence, we constructed single gene insertion-deletion strains in the wild type UPEC background (CFT073) by Lambda red recombination (42). We observed that CFT073 Δ*ihfB::kan* (JM028) and CFT073 Δ*zapE::kan* (JM023) strains grew robustly at low plating density (10^4^ CFU ml^-1^), similar to the mini-Tn5 mutants, suggesting that the deletion strains do not establish a quiescent state under the conditions tested, in contrast to the wild type strain (CFT073) (Fig. 1A).

Next, we tested if expression plasmids containing *ihfB* and *zapE* could restore quiescence (pBAD- *ihfB* and pBAD-*zapE*, respectively) of CFT073 Δ*ihfB::kan* (JM028) and CFT073 Δ*zapE::kan* (JM023) strains grown on glucose minimal agar at low plating density (10^4^ CFU ml^-1^). We observed that both deletion strains reverted to the quiescent phenotype when complemented by pBAD-*ihfB* and pBAD-*zapE*, respectively (Fig. 1B). In control assays, the vector (pBAD24) alone was unable to restore quiescence of either CFT073 Δ*ihfB::kan* (JM028) or CFT073 Δ*zapE::kan* (JM023) strains (Fig. 1B).

Next, we tested if known proliferants could prevent quiescence and promote proliferation of CFT073 Δ*ihfB::kan* (JM028) and CFT073 Δ*zapE::kan* (JM023) strains complemented with expression plasmids. Proliferant cues that were previously identified include peptidoglycan (PG) tetra-and pentapeptides and the amino acids L-lysine (1 mM) and L-methionine (1 mM) (28, 29). PG and the amino acids L-lysine and L-methionine (1 mM), stimulated proliferation of complemented strains (Fig. S1B).

### Antibiotic tolerance of quiescent *E. coli*

To test if quiescent *E. coli* cells are more tolerant to antibiotics than actively growing cells, we compared their survivability after exposure to ampicillin. First, we identified the minimum inhibitory concentration (MIC) of ampicillin for wild type UPEC (CFT073) and K-12 (MG1655) strains of *E. coli* cultured in liquid LB broth and found it to be similar (25 μg ml^-1^) (Fig. S2A). This indicates that CFT073 cells, which grow robustly in LB medium, and K-12 cells have similar antibiotic susceptibilities during growth in LB. To determine if quiescent cells are tolerant to ampicillin, we stimulated the quiescent state by adding UPEC cells to glucose minimal agar at low cell density. After 3 h at 37 °C, the cells were transferred to plates containing ampicillin (12.5 μg ml^-1^) for 20 h, and then cells were harvested and enumerated by viable plate counts on LB agar. We observed that under the conditions that induce the quiescent state, 3.6% of UPEC cells remained viable (Fig. 2A). In contrast, 0.01% of similarly cultured MG1655 cells were viable after ampicillin exposure. Notably, inoculation of CFT073 at a higher cell density, which prevents quiescence and promotes proliferation, reduced antibiotic tolerance by 1-log (Fig. S2B).

**Figure 2.**
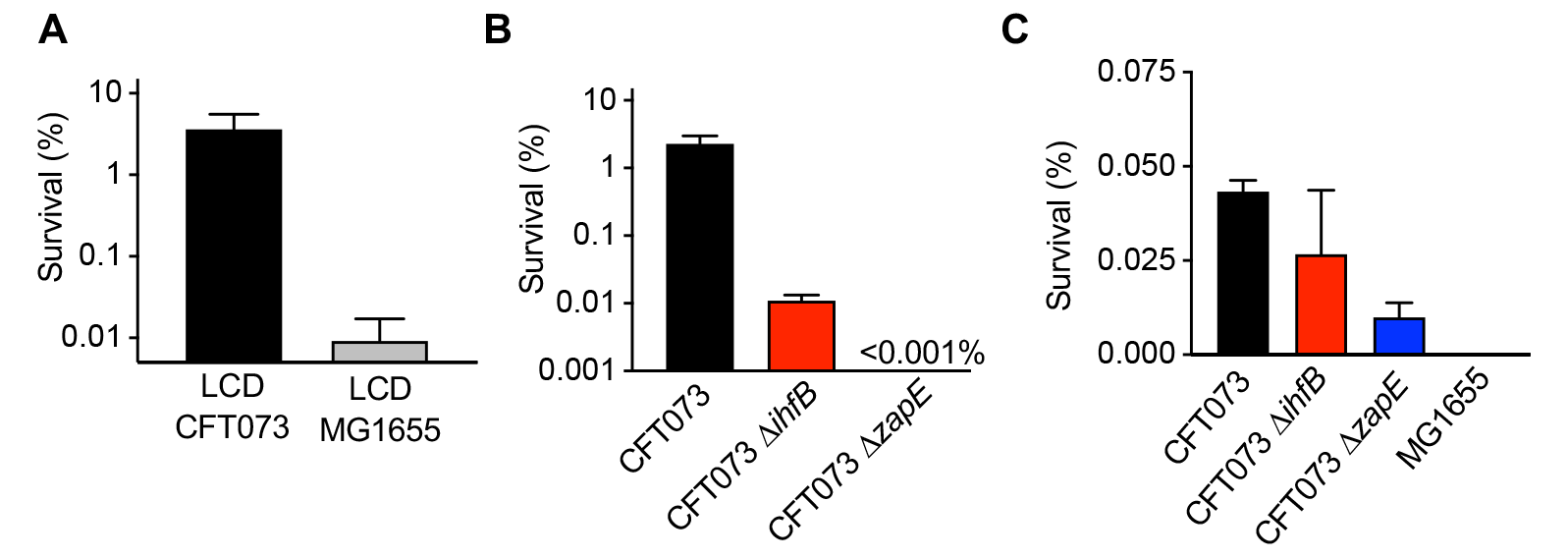
Quiescent UPEC CFT073 is antibiotic tolerant. (A) Antibiotic tolerance assay of CFT073 and MG1655 on ampicillin (12.5 μg ml^-1^) at low cell density (LCD) (10^4^ CFU ml^-1^), as described in Materials and Methods. Briefly, cells (10^5^ CFU ml^-1^) were spotted onto a 0.22 μm filter on M9 0.2% glucose agar and incubated at 37 °C for 3 h to induce quiescence for CFT073, then transferred to M9 0.2% glucose agar containing ampicillin (12.5 μg ml^-1^) and incubated at 37 °C for 20 h. Viability and percent survival was determined by serial dilution and spread plating. (B) CFT073, CFT073 Δ*ihfB,* and CFT073 Δ*zapE* tolerance to ampicillin (12.5 μg ml^-1^) at 10^4^ CFU ml^-1^ was assayed as in (A) and described in Materials and Methods. (C) Persister cell assay after 24 h as described in Materials and Methods. Briefly, overnight cultures were diluted to an OD_600_ of 0.1 in 0.2% glucose minimal media and incubated for 24 h at 37 °C and viable plates counts were determined to calculate percent survival. Data in (A) and (B) is an average of at least four independent experiments represented as mean ± SEM. Data in (C) is an average of at least three independent experiments represented as mean ± SEM.

To test if deletion of *zapE* or *ihfB*, which prevents quiescence, also reduces antibiotic tolerance, we compared viable plate counts of cells plated at low density (10^5^ CFU ml^-1^) for 3 h and then exposed to ampicillin for 20 h, as described. We observed that deletion of *zapE* or *ihfB* reduces tolerance to ampicillin by greater than 2-logs (Fig. 2B). This suggests that IhfB and ZapE are important for the antibiotic-tolerant quiescent state.

Antibiotic tolerance may also be detected in persister cell assays, which allows for quantitation of survivors following antibiotic exposure. In addition to establishing a quiescent state at low cell density on glucose minimal agar, cultures of wild type UPEC cells (CFT073) also display a high level of survivors in persister cell assays (29). To detect survivors, and test if deletion of *zapE* or *ihfB* reduces the level of survivors, we inoculated cultures with wild type (CFT073), Δ*zapE*, and Δ*ihfB* cells in glucose M9 minimal medium to an OD600 of 0.1 (approximately 10^8^ cells ml^-1^) in the absence and presence of ampicillin (100 μg ml^-1^), and then incubated the cultures at 37 °C and shaking. After 24 h, we quantitated the percent survivors in the ampicillin-containing cultures. We observed that 0.04 ± 0.003 % of wild type cells (CFT073) survive 24 h with ampicillin compared to 7.23 x 10^-5^ % for *E. coli* MG1655 (Fig. 2C), and deletion of either *ihfB* or *zapE* reduces the percentage of survivors to 0.025 ± 0.017% and 0.001 ± 0.003 %, respectively (Fig. 2C). Together, our results suggest that *ihfB* and *zapE* are essential for quiescence and important for antibiotic tolerance and survival with ampicillin.

### IHF and metabolic regulation of quiescence

To characterize the contribution of *ihfB* for quiescence, we evaluated the known and predicted genes in the IHF regulon. IhfB and IhfA coassemble into the IHF heterodimer, which constitutes the active regulatory complex to activate or repress transcription of target genes (30). If IhfB acts as part of the IHF complex to regulate genes important for quiescence, then deletion of *ihfA* should also prevent quiescence. To test this, we deleted *ihfA* by Lambda red recombination (CFT073 Δ*ihfA::kan*) and observed that CFT073 Δ*ihfA::kan* cells (JM068) grew robustly at low plating density (10^4^ CFU ml^-1^) on glucose minimal agar (Fig. 3A), similar to the non-quiescent growth observed after deletion of *ihfB* (Fig. 1A). However, in contrast to CFT073 Δ*ihfB::kan* (JM068), CFT073 Δ*ihfA::kan* (JM068) is not complemented by a plasmid containing *ihfB* (pBAD-*ihfB*) (Fig. 3A). Together, these results suggest that both *ihfA* and *ihfB* play a role in antibiotic-tolerant quiescence and implicate transcriptional regulation by IHF as a regulatory factor.

**Figure 3.**
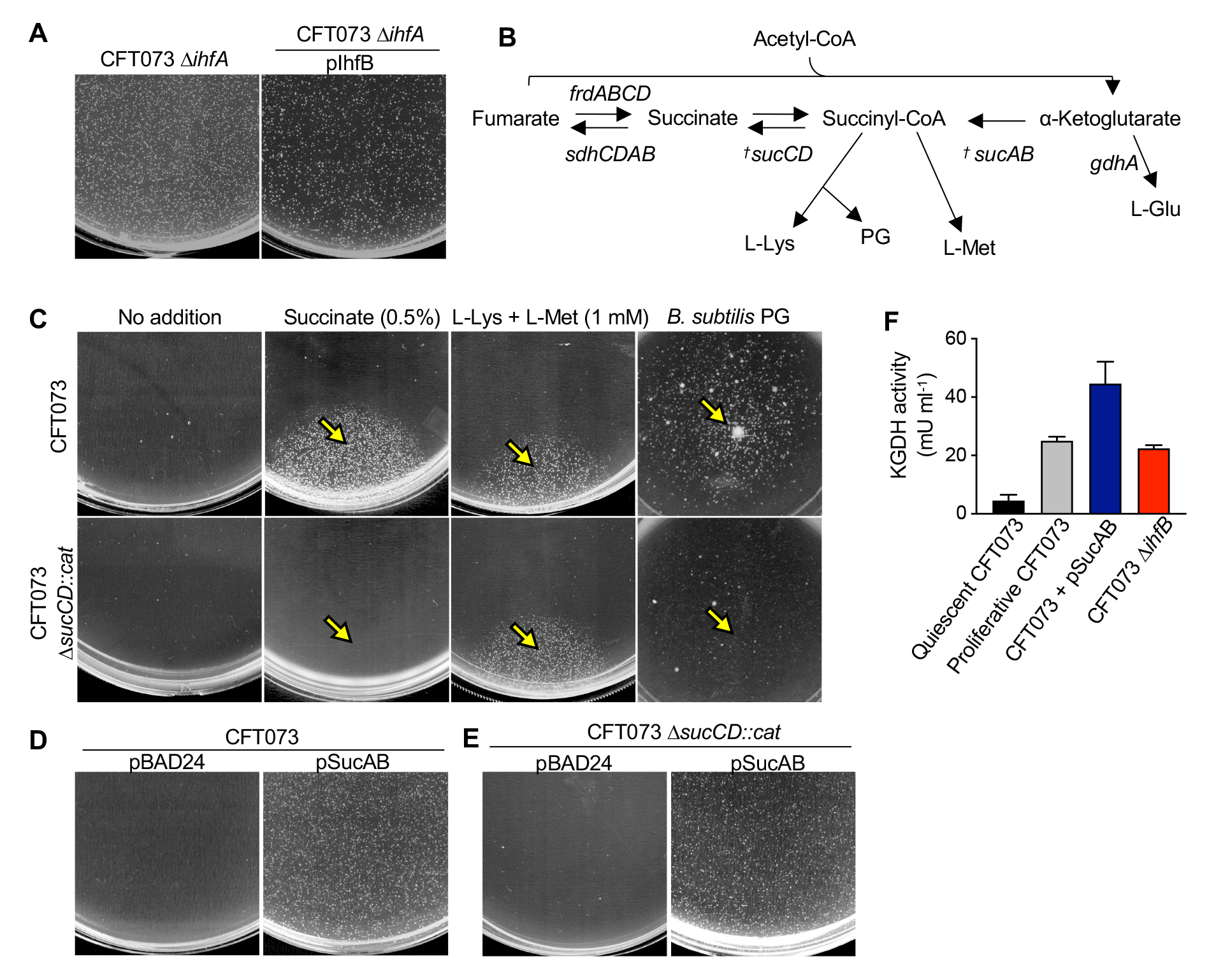
Importance of succinyl-CoA in UPEC metabolically inactive quiescence. (A) CFT073 Δ*ihfA* and CFT073 Δ*ihfA* pIhfB plated at 10^4^ CFU ml^-1^ on 0.2% glucose minimal agar. Plates containing CFT073 Δ*ihfA* pIhfB were supplemented with 0.1% arabinose and 50 μg ml^-1^ ampicillin and were incubated at 37 °C for 24 hrs. (B) Abbreviated TCA cycle showing α- Ketoglutarate conversion into succinyl-CoA and subsequent conversion into succinate and fumarate. Succinyl-CoA is utilized for L-Lys, L-Met, and peptidoglycan biosynthesis. *^†^* refers to genes regulated by IHF (37, 38). (C) CFT073 and CFT073 Δ*sucCD* and were plated at 10^4^ CFU ml^-1^ on 0.2% glucose minimal agar and then challenged with 5 μl of succinate (0.5%), a combination of L-Lys and L-Met (1 mM), or mutanolysin digested *B. subtilis* peptidoglycan (PG) as described in Materials and Methods. Yellow arrows represent where stimulants were added. Plates were incubated at 37 °C for 24 hrs. (D) CFT073 pBAD24 or CFT073 pSucAB were plated at 10^4^ CFU ml^-1^ on 0.2% glucose minimal agar supplemented with 0.1% arabinose and 50 μg ml^-^ _1_ ampicillin. Plates were incubated at 37 °C for 24 hrs. (E) CFT073 Δ*sucCD* pBAD24 and CFT073 Δ*sucCD* pSucAB were plated at 10^4^ CFU ml^-1^ on 0.2% glucose minimal agar supplemented with 0.1% arabinose and 50 μg ml^-1^ ampicillin. Plates were incubated at 37 °C for 24 hrs. (F) α- Ketoglutarate dehydrogenase assay was performed as described in materials and methods on quiescent and proliferative CFT073, CFT073 pSucAB, or CFT073 Δ*ihfB* using the KGDH assay kit (Sigma-Aldrich). Pictures in (A), (C), (D), and (E) are representative of at least three independent experiments. Data in (F) is an average of at least three independent experiments represented as mean ± SEM.

The IHF regulon includes metabolic enzymes of the tricarboxylic acid cycle (TCA), including genes important for regulating the levels of succinyl-CoA (*sucA* and *sucB*) and succinate (*sucC* and *sucD*) (Fig. 3B). A gene that was previously implicated in quiescence, *sdhA*, is also critical for converting succinate to fumarate in the TCA cycle (29) (Fig. 3B). Deletion of *sdhA* would likely lead to an accumulation of succinate (43). To directly test if succinate reverses quiescence, we performed the quiescence assay on wild type cells (CFT073) plated at low density on glucose minimal agar and compared the addition of succinate to known proliferant cues PG and amino acids L-lysine (1 mM) and L-methionine (1 mM). We observed that succinate (0.5%) caused a robust zone of growth similar to other known stimulants (Fig. 3C).

Succinate is generated from succinyl-CoA by succinyl-CoA synthetase, a bidirectional TCA cycle enzyme complex that regulates succinate/succinyl-CoA pools and is encoded by *sucC* (beta subunit) and *sucD* (alpha subunit) (Fig. 3B). To determine if the succinate added to culture medium and shown to reverse quiescence requires conversion to succinyl-CoA, we deleted *sucC* and *sucD* from the *suc* operon and then determined if succinate overrides quiescence in the deletion strain as in the wild type strain. We observed that CFT073 Δ*sucCD::cat* (JM071) cells were quiescent on glucose minimal agar, similar to wild type cells (CFT073); however, they failed to proliferate with addition of succinate (Fig. 3C). In contrast, CFT073 Δ*sucCD::cat* (JM071) cells were stimulated to proliferate with amino acids (L-lysine and L-methionine) and PG (Fig. 3B), indicating that they are not defective for exiting the quiescent state.

IHF was reported to repress *sucA* and *sucB* expression by 3-4 fold (37, 38). Together with lipoamide dehydrogenase, SucA and SucB form the α-ketoglutarate dehydrogenase (KGDH) complex responsible for conversion of α-ketoglutarate to succinyl-CoA. Therefore, deletion of *ihfB* could relieve transcriptional repression of the *suc* operon (*sucA, sucB, sucC, sucD*), thus increasing succinate pools in the cell and overall flux through the TCA cycle. To test if overexpression of *sucA* and *sucB* is sufficient to override quiescence in UPEC cells, we constructed a *sucAB* expression plasmid and induced expression of both genes in the operon with arabinose in wild type UPEC cells (CFT073). Cells containing pSucAB and cultured at low cell density on glucose minimal agar, supplemented with 0.1% arabinose and ampicillin (50 μg ml^-1^) were non-quiescent, in contrast to cells containing the control vector (pBAD24), which were quiescent (Fig 3D). The requirement of *sucCD* for reversal of quiescence by succinate suggests that succinyl-CoA is the critical metabolite for regulation of quiescence. To test this, we transformed CFT073 Δ*sucCD:cat* (JM071) cells with pSucAB to prevent subsequent conversion to succinate. We observed that the plasmid containing *sucAB*, encoding KGDH to convert α-ketoglutarate to succinyl-CoA, also reversed quiescence in cells deleted for *sucCD* (Fig. 3E). This result is consistent with succinyl-CoA functioning as a critical metabolite for overcoming quiescence.

Lastly, we directly measured KGDH activity in quiescent and proliferating UPEC cells. We observed that quiescent CFT073 had very low KGDH activity compared to proliferating cells (Fig. 3F). Cells deleted for *ihfB* had KGDH activity similar to proliferating cells, and, as expected, cells containing pSucAB had the highest level of activity measured in the assay (Fig. 3F). This suggests that quiescent UPEC cells have lower overall KGDH activity and reduced metabolism. Together, these results suggest that succinyl-CoA availability both prevents and reverses quiescence in UPEC. Thus, quiescent cells are likely limited for succinyl-CoA. Notably, succinyl-CoA is required for biosynthesis of L-lysine, L-methionine, and peptidoglycan precursors (diaminopimelate) (Fig. 3B) (44, 45).

### The cell division proteins ZapE and FtsN engage SDH complexes

ZapE has been reported to be important for cell division in *E. coli* exposed to various stress conditions, including elevated temperature and low oxygen environments (31, 32). ZapE binds directly to FtsZ and phospholipids (32), and also interacts with late-stage cell division proteins, including FtsN, FtsI, FtsQ and FtsL in bacterial two hybrid assays performed in a K-12 strain (31). Notably, quiescent UPEC cells have been reported to be filamentous after extended incubation on glucose minimal agar, suggesting that cell division is impaired or blocked when cells are in the quiescent state (28). First, we examined the morphology of wild type cells (CFT073) and cells deleted for *zapE*. After incubating cells plated at low cell density on glucose minimal agar for 24 h, the conditions that support quiescence, wild type CFT073 cells were filamentous, with an average cell length of 5.2 ± 4.1 μm (n=200) (Fig. 4A), and we observed 33% of cells longer than 5 μm, in agreement with previous reports (28). In contrast, cells deleted for *zapE*, which are unable able to enter the quiescent state, were shorter, with an average cell length of 2.9 ± 2.1 μm (n=200) (Fig. 4A), and only 6.5% of cells were longer than 5 μm. This suggests that quiescent cells are non-proliferative and mildly filamentous, and deletion of *zapE* restores proliferative growth, likely allowing cells to complete division cycles thus producing overall shorter cells. Notably, as deletion of *zapE* was previously reported to induce filamentation in *E. coli* K-12 strains (31, 32), we also visualized wild type and *zapE* deletion strains cultured at high inoculation density on LB agar for 6 h. We observed a small percentage (3%) of filamentous cells present in cultures of CFT073 Δ*zapE:kan* strain (JM023); however, wild type CFT073 cells were non-filamentous (Fig. S3A). Finally, in wild type and *zapE* deletion strains, cells expressing *zapE* from a plasmid were also non-filamentous (Fig. S3B).

**Figure 4.**
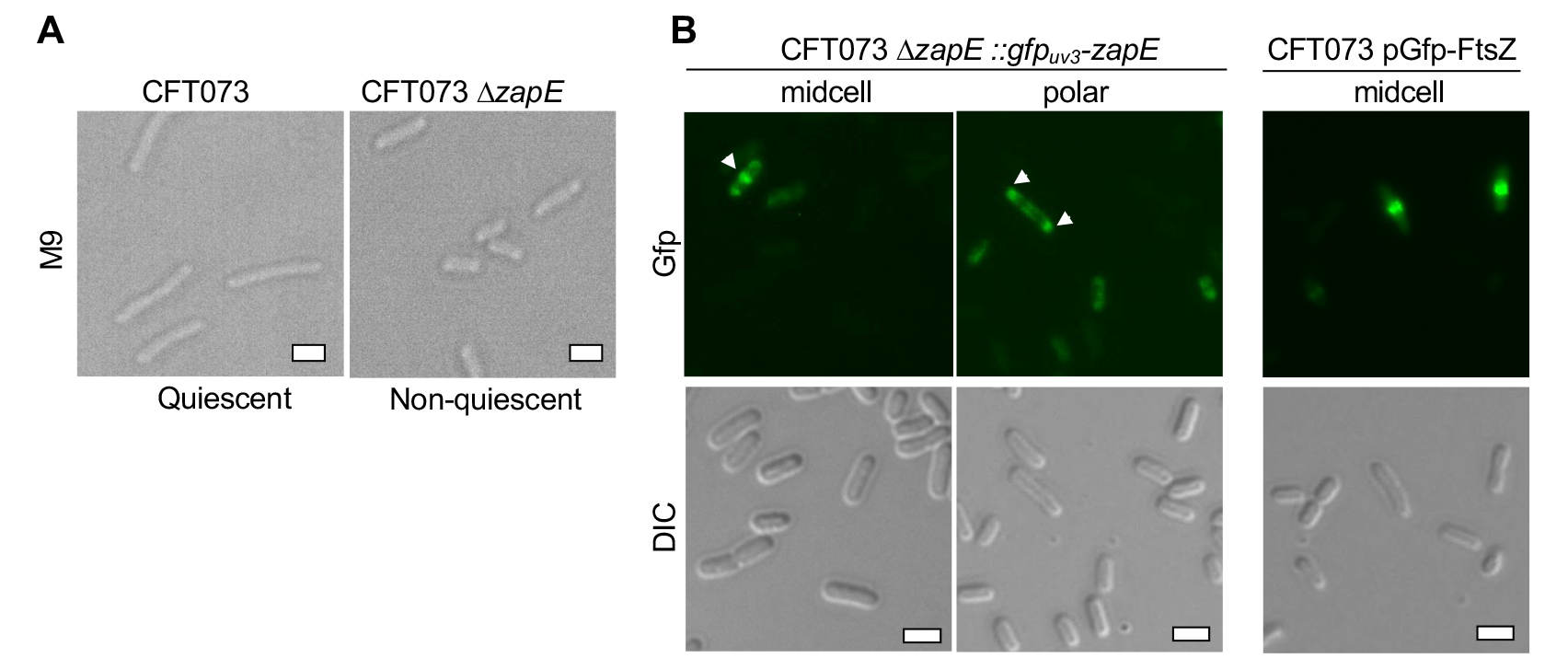
Morphology of CFT073 *zapE* deletion and overexpression, and localization of Gfp_uv3_-ZapE. (A) Differential interference contrast (DIC) microscopy of CFT073 or CFT073 Δ*zapE* strains inoculated at 10^6^ CFU ml^-1^ on glucose (0.2%) minimal agar and then incubated for 24 h at 37 °C before being harvested. (B) Epifluorescence microscopy of CFT073 expressing Gfp-FtsZ from pBAD24 or CFT073 Δ*zapE*::*gfp-zapE_uv3_*. Cells expressing Gfp-FtsZ were grown for 6 h at 30 °C on LB agar supplemented with 100 μg ml^-1^ ampicillin and 0.01% arabinose and cells expressing Gfp_uv3_-ZapE were grown on LB agar. Scale bars in (A), (B), and (C) are 2 μm.

ZapE was previously reported to localize to late stage division rings in an *E. coli* K-12 strain when expressed as a fluorescent fusion protein (31). To test if ZapE also localizes to the division site in UPEC cells that are actively dividing, we expressed *gfp-zapE* from the native locus in the chromosome. Briefly, we used λ red recombination to replace the antibiotic resistance cassette of CFT073 Δ*zapE* cells with *gfp_uv3_-zapE*, as described in Materials and Methods (42, 46). We cultured cells expressing Gfp-ZapE on LB agar for 6 h and visualized cells by fluorescence microscopy. We observed midcell localization of Gfp-ZapE in 5.5% of cells, consistent with Z-ring recruitment. We also observed polar localization of Gfp-ZapE in 6% of cells (Fig. 4B), suggesting that Gfp-ZapE may remain at the site for some time after division completes. Finally, as a reference for Z-ring assembly, we also visualized localization of Gfp-FtsZ in wild type CFT073 cells that are actively dividing and observed robust Z-rings (Fig. 4B).

We previously determined that low succinyl-CoA and reduced metabolism is critical for quiescence, therefore, we investigated if ZapE has a role in modulating metabolism. We tested if the non-quiescent phenotype in the *zapE* deletion strain requires the conversion of succinate to succinyl-CoA. We deleted *sucCD* from the CFT073 Δ*zapE* strain to construct CFT073 Δ*zapE:kan* Δ*sucCD:cat* (JM212) and tested if the multiple deletion strain remained non-quiescent. We observed that the additional deletion of *sucCD* restored the quiescent phenotype of cells deleted for *zapE* (Fig. 5A). Moreover, quiescent CFT073 Δ*zapE:kan* Δ*sucCD:cat* cells proliferate in response to L-lysine and L-methionine, but not in response to succinate, similar to Δ*sucCD* cells (Fig. 5A) (Fig. 3C). This result suggests that ZapE may regulate quiescence through engaging TCA cycle enzymes in a manner that limits succinate or succinyl-CoA.

**Figure 5.**
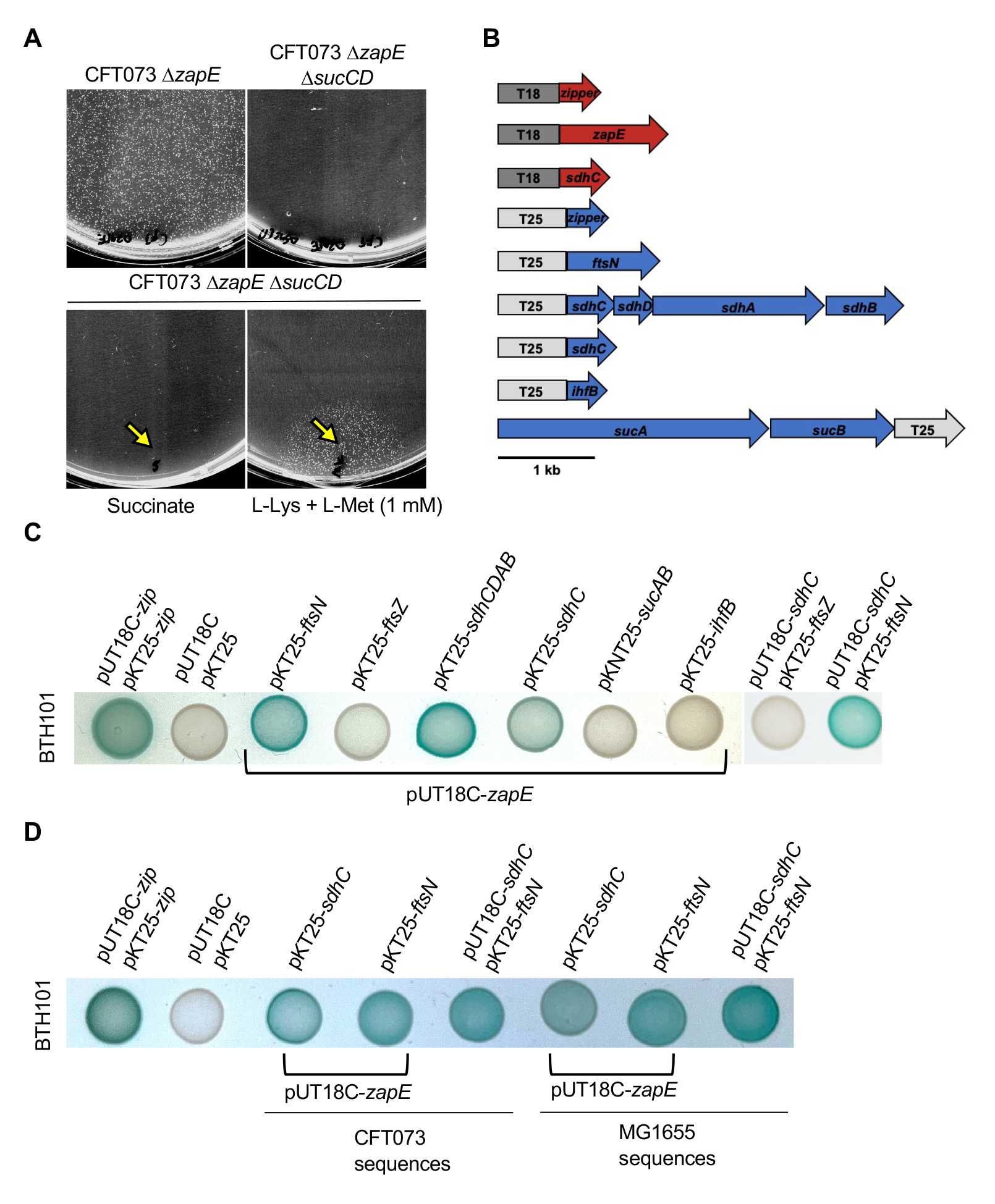
ZapE requires *sucCD* for its role in blocking UPEC division and interacts with components of the cell division machinery and the TCA cycle. (A) CFT073 Δ*zapE* and CFT073 Δ*zapE* Δ*sucCD* were plated at 10^4^ CFU ml^-1^ on 0.2% glucose minimal agar. Where indicated, CFT073 Δ*zapE* Δ*sucCD* cells were challenged with 5 μl of succinate (0.5%), or a combination of L-Lys and L-Met (1 mM). Plates were incubated at 37 °C for 24 hrs. Yellow arrows represent where stimulants were added. (B) Diagram of gene plasmid construction of plasmids (pUT18C, pKT25, or pKNT25) cloned for the BACTH assay. (C and D) BTH101 co-expressing plasmids for X-gal assay plated on LB agar containing ampicillin (100 μg ml^-1^), kanamycin (50 μg ml^-1^), IPTG (0.5 mM), and X-gal (40 μg ml^-1^). Plates were incubated for 24 hours at 30 °C. Pictures in (A), (C), and (D) are representative of at least three independent experiments.

Finally, since the bacterial two hybrid assay has been used to successfully detect interactions between ZapE and other cell division proteins (33), we tested if ZapE interacts with TCA cycle enzymes that regulate succinate and/or succinyl-CoA. We constructed T18 and T25 adenylate cyclase fusion proteins to detect interactions between ZapE and components of IHF, α- ketoglutarate dehydrogenase (KGDH), succinate dehydrogenase (SDH), and FtsN. We fused the N-terminus of the *zapE* gene to T18 using vector pUT18C (Fig. 5B). All other genes tested were cloned with T25 fused to the N-terminus using vector pKT25, or with the T25 fused to the C-terminus in the case of *sucAB* using the vector pKNT25 (Fig. 5B). The vectors were co-transformed into *E. coli* strain BTH101 which lacks the *cya* gene encoding for adenylate cyclase and evaluated potential interactions via plating on nutrient agar plates containing X-gal (5-Bromo-4-chloro-3-indolyl β-D-galactopyranoside). In control assays, we observed that, when co-expressed, T18 and T25 fused to a leucine zipper (*zip*) known to interact lead to blue colonies, indicating interaction, while a white colony appearance was observed when T18 and T25 were co-expressed alone (Fig. 5C). Then, we evaluated a known interaction between ZapE and FtsN (31). As expected, we observed a robust colorimetric change when co-expressing T18-ZapE with T25- FtsN, as previously reported (Fig. 5C) (31). The SDH complex is encoded by the *sdhCDAB* operon, and *sdhA* is essential for UPEC quiescence (29). Therefore, we evaluated an interaction between T18-ZapE and T25-SdhC alone, and also engineered the plasmid to contain the full *sdhCDAB* operon on pKT25. We observed a colorimetric change in cells expressing T18-ZapE and T25- SdhC alone, which increased in intensity when the full *sdhCDAB* operon was included in the expression plasmid (Fig. 5B and 5C). We detected no interactions between T18-ZapE and T25- IhfB, SucAB-T25, SucA-T25, or SucB-T25 (Fig. 5C and S3C). By bacterial two hybrid, ZapE interacts with both FtsN and SdhC, therefore we tested if SdhC interacts directly with FtsN. In cells expressing T18-SdhC and T25-FtsN, we detected a large colorimetric change suggestive of a direct interaction. Interestingly, we did not detect any interaction between FtsZ and SdhC, or between FtsZ and ZapE in this assay, although ZapE is known to bind to FtsZ polymers (31, 32). We also performed the Miller assay to quantitatively compare several of the interactions and observed trends similar to the result on X-Gal plates (Fig. S3D).

Cell division proteins are not known to engage TCA cycle enzymes in *E. coli,* therefore we compared amino acids sequences of ZapE, FtsN, and SdhC in several strains, including CFT073, the K-12 strain MG1655, and Nissle 1917, a probiotic strain with genetic similarity to CFT073 (47). While FtsN is highly conserved, nine amino acid substitutions have been incorporated into ZapE in CFT073 compared to the MG1655 strain. In addition, SdhC in CFT073 and Nissle 1917 strains are annotated to contain five additional amino acids at the N-terminus, which are not present in MG1655 SdhC (Fig. S4A). ZapE has the most amino acid variation, therefore we compared ZapE sequences across bacterial families and found that ZapE is fairly well conserved at the amino acid sequence level in *Enterobacteriaceae* but not well conserved in other organisms with ZapE. (Fig. S4B). To determine if ZapE from K-12 interacts with SdhC and FtsN, we repeated the bacterial two hybrid assay with K-12-derived fusion proteins and again observed that ZapE interacts with both SdhC and FtsN (Fig. 5D). Additionally, we observed that SdhC and FtsN from *E. coli* K-12 also interact (Fig. 5D). These results demonstrate that interactions between cell division proteins and SDH complexes, which are present at the inner membrane, are likely conserved in *E. coli* and other *Enterobacteriaceae*, including in organisms that are not known to enter a quiescent state.

### Metabolic flux regulates UPEC antibiotic tolerance

We determined that supplementing cells exogenously with succinate or increasing intracellular levels of succinyl-CoA by expressing *sucAB* from a plasmid reversed the quiescent state (Fig. 3C and 3D). As antibiotic tolerance appears to be strongly associated with quiescence, we tested if succinate increases antibiotic susceptibility of UPEC on glucose minimal agar. Briefly, we added cells to glucose minimal agar at low cell density to induce the quiescent state. After 3 h at 37 °C, the cells were transferred to glucose minimal plates containing ampicillin (12.5 μg ml^-1^), with and without succinate (0.5%) for 20 h, and then cells were harvested and enumerated by viable plate counts on LB agar. We observed that under the conditions that induce the quiescent state, 1% of UPEC cells remained viable (Fig. 6A), similar to our previous result (Fig. 2A). However, when wild type, quiescent cells were exposed to succinate and ampicillin for 20 h, the majority of cells were killed, and viability was less than 0.001% (Fig. 6A)

**Figure 6.**
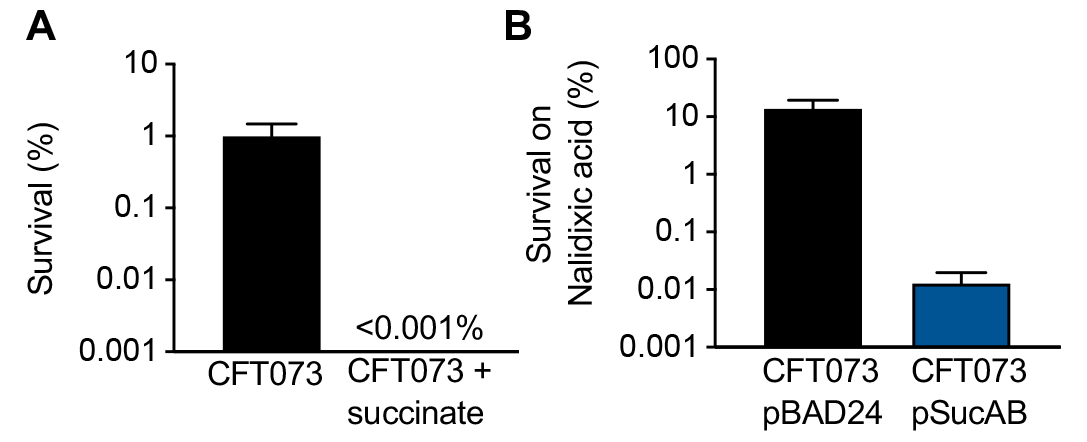
Metabolic flux regulates UPEC antibiotic tolerance. (A) Antibiotic tolerance assay of CFT073 and CFT073 supplemented with succinate (0.5%), on ampicillin (12.5 μg ml^-1^) as described in Materials and Methods. (B) Antibiotic tolerance assay of CFT073 pBAD24 and CFT073 pSucAB on nalidixic acid (7.5 μg ml^-1^) as described in Materials and Methods. 0.2% glucose minimal agar plates were supplemented with 0.1% arabinose and 50 μg ml^-1^ ampicillin.

We also tested if overexpression of SucAB promotes increased antibiotic susceptibility in the antibiotic tolerance assay. We compared sensitivity of wild type cells containing pSucAB or the control vector (pBAD24) to nalidixic acid, since ampicillin was used for plasmid selection. We observed that under the conditions that induce the quiescent state, 13.8% of UPEC cells containing the control vector remained viable after incubation with nalidixic acid (7.5 μg ml^-1^) for 20 h (Fig. 6B). However, cells containing the pSucAB expression plasmid, were effectively killed by nalidixic acid and viability was 0.01% (Fig. 6B). Together, our results demonstrate that quiescent cells are antibiotic-tolerant, and tolerance is reversed by succinate or by increasing KGDH activity.

## Discussion

Here, we describe a fundamental strategy that UPEC cells use to regulate quiescence and antibiotic tolerance through the action of the IHF transcriptional regulator complex and ZapE, a cell division protein that also interacts with succinate dehydrogenase (Fig. 1, 5C). We show that the UPEC quiescent state is critical for antibiotic tolerance and is modulated by succinate levels (Fig. 2 and 6). Limitation of succinyl-CoA in the TCA cycle is critical for maintenance of UPEC quiescence and quiescent cells have slowed cell metabolism (Fig. 3). As IHF is known to repress *sucA* and *sucB* expression (37, 38), it follows that this may be the primary mechanism by which IHF suppresses proliferation and promotes quiescence in this UPEC strain. Lastly, we show that ZapE, which interacts with FtsZ and FtsN of the cell division machinery, also interacts with succinate dehydrogenase of the TCA cycle suggesting it may act as link between division and metabolism in *E. coli* (Fig. 5).

UPEC has been shown to form QIRs in a murine model system where they may lie dormant for extended periods of time, maintain antibiotic tolerance, and potentially contribute to UTI recurrence (19, 24, 48, 49). We show that non-proliferating, quiescent UPEC is tolerant to the bactericidal antibiotic ampicillin in our in vitro quiescence assay (Figs. 2A and B). Additionally, we observed that UPEC strain CFT073 generates a high level of persister cells in contrast to *E. coli* strain MG1655, which is in agreement with a previous report (29). We also observed that deletion of *zapE* or *ihfB* reduced the amount of persisters under the conditions tested, supporting that reduced growth and metabolism is important for persistence (Fig. 2C). IHF has recently been reported to be important for persister cell formation by rerouting TCA cycle metabolism through upregulation of *aceA* in the glyoxylate bypass and downregulation of *icd* in the TCA cycle (39). Bactericidal antibiotics such as ampicillin are known to increase metabolic activity, and reduced cell metabolism increases tolerance to bactericidal antibiotics (13). For example, bacterial cells embedded deep within biofilms have marked reduced cell metabolism which prevents the effects of bactericidal antibiotics (50, 51). Consistent with this, we observed quiescent CFT073 cells also have low KGDH activity indicating reduced cell metabolism (Fig. 3F). Addition of succinate or overexpression of SucAB (KGDH components) resuscitated quiescent cells and caused increased antibiotic susceptibility indicating that cell metabolism is critical for quiescence and antibiotic tolerance in UPEC (Fig. 6).

Together, our results suggest that succinyl-CoA is likely limited while cells are in the quiescent state as addition of succinate stimulates proliferation, although not in the *sucCD* deletion strain, and cells expressing SucAB (KGDH) are non-quiescent (Figs. 3C, D, and E). Previously, *gdhA* and *sdhA* were identified by a mini transposon screen as critical for quiescence in UPEC (29). Deletion of either of these genes would lead to accumulation of succinyl-CoA, which is consistent with our findings. Interestingly, an UPEC *sdhB* mutant was found to be defective for colonization in a mouse infection model but not defective in vitro for growth in urine (52). Consequently, it would be interesting to determine if strategies to modulate succinyl-CoA levels in UPEC would prevent infection and/or antibiotic tolerance in an in vivo model.

Both subunits of the regulatory IHF complex, IhfA and IhfB, are critical for the antibiotic tolerant, quiescent state in our in vitro system with UPEC strain CFT073 (Figs. 1A and 3A). Interestingly, the IHF complex has previously been shown to be critical for UPEC establishment of UTIs and stabilizes pathogenicity island I in UPEC strain 536 (36, 53). IHF also appears to be critical for establishment of intracellular bacterial communities (IBCs) in a murine model (54). The IHF complex is reported to regulate more than 100 genes in *E. coli* and repress *sucAB* expression by 3- 4 fold (37, 38). However, with the complexity and size of the IHF regulon, it is difficult to conclusively evaluate its contribution to quiescence.

ZapE was previously reported to be important for cell division during periods of cell stress, under oxygen-limited growth (31, 32). We observed that ZapE is essential for quiescence and interacts with the cell division machinery and with TCA cycle components, indicating it may act as a link to coordinate cell division with metabolism (Figs. 1 and 5). Pyruvate levels regulate transcription of the *fts* operon through the action of PdhR (55, 56); however, no direct interactions between cell division proteins and TCA cycle enzymes have been reported in *E. coli*. However, in *Bacillus subtilis*, the E1α subunit of pyruvate dehydrogenase interacts with the nucleoid under high nutrient conditions and may contribute to FtsZ placement at midcell during nucleoid segregation (57). Another *E. coli* protein that coordinates metabolism with division is the inner membrane protein OpgH, which acts as a UDP-glucose sensor to antagonize FtsZ polymerization when high levels of UDP-glucose are present (58). OpgH may act as a proxy for nutritional status to ensure cells coordinate division with metabolism. Here, our work demonstrates direct interactions among ZapE, FtsN and SdhC that are conserved in *E. coli.* However, the mechanism by which ZapE facilitates quiescence is still not clear. ZapE may alter SDH activity directly, a hypothesis that is consistent with the role of succinyl-CoA in the quiescence phenotype described in this study. However, as ZapE is known to interact with FtsZ, FtsN, and other late-stage cell division proteins (31, 32),it is also possible that ZapE halts division progression through a direct interaction with cell division proteins independent of its relationship with SDH complexes. Notably, we observed that *E. coli* K-12 ZapE interacts with K-12 SdhC and FtsN in the bacterial two hybrid assay (Fig. 5D), indicating that the link between division and SDH is conserved in *E. coli* strains that are non-pathogenic and do not enter a quiescent state. Future studies will focus on elucidating the regulation between TCA cycle flux and division progression.

## Materials and methods

### Bacterial strains and growth conditions

*E. coli* strains and plasmids used in this study are included in table 1. Primers used for gene cloning and gene deletions are shown in table 2. Plasmids pUT18C-*zapE* (CFT073), pUT18C-*zapE* (MG1655), pUT18C-*sdhC* (CFT073), pUT18C-*sdhC* (MG1655), pKT25-*ftsN* (CFT073), pKT25- *ftsN* (MG1655) pKT25-*ftsZ*, pKT25-*sdhCDAB* (CFT073), pKT25-*sdhC* (CFT073), pKT25-*sdhC* (MG1655), pKNT25-*sucAB* (CFT073), pKNT25-*sucA* (CFT073), and pKNT25-*sucB* (CFT073) were constructed by amplifying genes from *E. coli* CFT073 or MG1655 via whole colony PCR using primers PstIzapEfor and EcoRIZapErev(for pUT18C-*zapE* (CFT073) or pUT18C-*zapE* (MG1655)), sdhCpUT18Cfor and sdhCpUT18Crev (for pUT18C-*sdhC* (CFT073)), K12sdhCpUT18Cfor and sdhCpUT18Crev (for pUT18C-*sdhC* (MG1655)) PstIftsNfor and EcoRIFtsNrev(for pKT25-*ftsN* (CFT073) or pKT25-*ftsN* (MG1655)), PstIFtsZfor and BamHIFtsZrev(for pKT25-*ftsZ*), sdhCDABPstIF and sdhCDABEcoRIR(for pKT25-*sdhCDAB* (CFT073)), sdhCDABPstIF and SdhCEcoRIrev(for pKT25-*sdhC* (CFT073)), K12sdhCpKT25 and sdhCpUT18Crev (for pKT25-*sdhC* (MG1655)), HindIIIsucABfor and sucABEcoRIrev(for pKNT25-*sucAB* (CFT073)), HindIIIsucABfor and EcoRIsucArev (for pKNT25-*sucA* (CFT073)), and HindIIIsucBfor and sucABEcoRIrev (for pKNT25-*sucB* (CFT073)) (table 2). The resulting PCR products were double digested with PstI(HiFi) and EcoRI(HiFi) restriction enzymes (or HindIII(HiFi) and EcoRI(HiFI) for pKNT25-*sucAB,* pKNT25-*sucB,* and pKNT25-*sucA,* or PstI(HiFI) and BamHI(HiFi) for pKT25-*ftsZ*) and purified via gel purification. Digested PCR products were ligated into PstI/EcoRI digested pUT18C or pKT25 (or PstI/BamHI digested pKT25 for pKT25-*ftsZ* or HindIII/EcoRI digested pKNT25 for pKNT25-*sucAB*, pKNT25-*sucB*, or pKNT25-*sucA*). Ligation reactions were transformed into XL10 gold ultracompetent cells (Agilent) and verified by sequencing. Plasmid mixtures of pUT18C-*zapE* (CFT073 or MG1655) or pUT18C-*sdhC* (CFT073 or MG1655) and each other individual BACTH plasmids to be tested were transformed into *E. coli* BTH101 strains and plated onto LB agar containing ampicillin (100 μg ml^-1^) and kanamycin (50 μg ml^-1^). These dual plasmid containing strains were then further analyzed via the X-Gal or Miller assay (see bacterial two-hybrid assay (BACTH) X-Gal and Miller assays).

**Table 1.** *E. coli* strains and plasmids.

| Strain or plasmid | Relevant Genotype | Source, reference, or construction |
| --- | --- | --- |
| **Strain** |  |  |
| CFT073 Str<sup>r</sup> | Spontaneous streptomycin<br>-resistant mutant of CFT073 | (29) |
| CFT073 | Original clinical isolate | (61) |
| MG1655 | LAM- *-rph1* | (62) |
| JM205 | CFT073 Str<sup>r</sup> *ihfB Tn5::kan* | This study |
| JM206 | CFT073 Str<sup>r</sup> *zapE Tn5::kan* | This study |
| JM023 | CFT073 Str<sup>r</sup> Δ*zapE::parE-kan* | λred recombination, this study |
| JM028 | CFT073 Str<sup>r</sup> Δ*ihfB::parE-kan* | λred recombination, this study |
| JM068 | CFT073 Str<sup>r</sup> Δ*ihfA::parE-kan* | λred recombination, this study |
| JM071 | CFT073 Str<sup>r</sup> Δ*sucCD::cat* | λred recombination, this study |
| JM120 | CFT073 Str<sup>r</sup> Δ*zapE::gfp<sub>uv3</sub>-zapE* | λred recombination, this study |
| JM212 | CFT073 Str<sup>r</sup> Δ*zapE::parE-kan*<br>Δ*sucCD::cat* | λred recombination, this study |
| PC358 | CFT073 Str<sup>r</sup> *sdhA Tn5::kan* | (29) |
| BTH101 | F-, *cya-99, araD139, galE15,*<br>*galK16, rpsL1 (Str r), hsdR2,*<br>*mcrA1, mcrB1* | (63) |
| **Plasmid** |  |  |
| pKD3 | *cat* | (42) |
| pKD46 | *amp* | (42) |
| pKD267 | *kan* | B. Wanner<sup>a</sup> |
| pBAD24 | *amp* | (64) |
| pBAD24-*zapE* | *amp P<sub>ara</sub>::zapE* | This study |
| pBAD24-*ihfB* | *amp P<sub>ara</sub>::ihfB* | This study |
| pBAD24-*sucAB* | *amp P<sub>ara</sub>::sucAB* | This study |
| pBAD24-*gfp<sub>uv3</sub>* | *amp P<sub>ara</sub>::gfp<sub>uv3</sub>* | Lab stock |
| pBAD24-*gfp<sub>uv3</sub>-ftsZ* | *amp P<sub>ara</sub>::gfp<sub>uv3</sub>-ftsZ* | (65) |
| pBAD24-*gfp<sub>uv3</sub>-zapE* | *amp P<sub>ara</sub>::gfp<sub>uv3</sub>-zapE* | This study |
| pUT18C | *amp* | (63) |
| pKT25 | *kan* | (63) |
| pKNT25 | *kan* | (63) |
| pUT18C-*zip* | *amp P<sub>lac</sub>::Leucine Zipper* | (63) |
| pKT25-*zip* | *kan P<sub>lac</sub>::Leucine Zipper* | (63) |
| pUT18C-*zapE* (CFT073) | *amp P<sub>lac</sub>::zapE* | This study |
| pUT18C-*zapE* (MG1655) | *amp P<sub>lac</sub>::zapE* | This study |
| pKNT25-*sucAB* (CFT073) | *kan P<sub>lac</sub>::sucAB* | This study |
| pKNT25-*sucA* (CFT073) | *kan P<sub>lac</sub>::sucAB* | This study |
| pKNT25-*sucB* (CFT073) | *kan P<sub>lac</sub>::sucAB* | This study |
| pKT25-*ftsN* (CFT073) | *kan P<sub>lac</sub>::ftsN* | This study |
| pKT25-*ftsN* (MG1655) | *kan P<sub>lac</sub>::ftsN* | This study |
| pKT25-*ftsZ* | *kan P<sub>lac</sub>::ftsZ* | This study |
| pKT25-*sdhCDAB* (CFT073) | *kan P<sub>lac</sub>::sdhCDAB* | This study |
| pKT25-*sdhC* (CFT073) | *kan P<sub>lac</sub>::sdhC* | This study |
| pKT25-*sdhC* (MG1655) | *kan P<sub>lac</sub>::sdhC* | This study |
| pUT18C-*sdhC* (CFT073) | *amp P<sub>lac</sub>::sdhC* | This study |
| pUT18C-*sdhC* (MG1655) | *amp P<sub>lac</sub>::sdhC* | This study |

<sup>a</sup>J. Teramoto, K. A. Datsenko, and B. L. Wanner, unpublished.

**Table 2.**
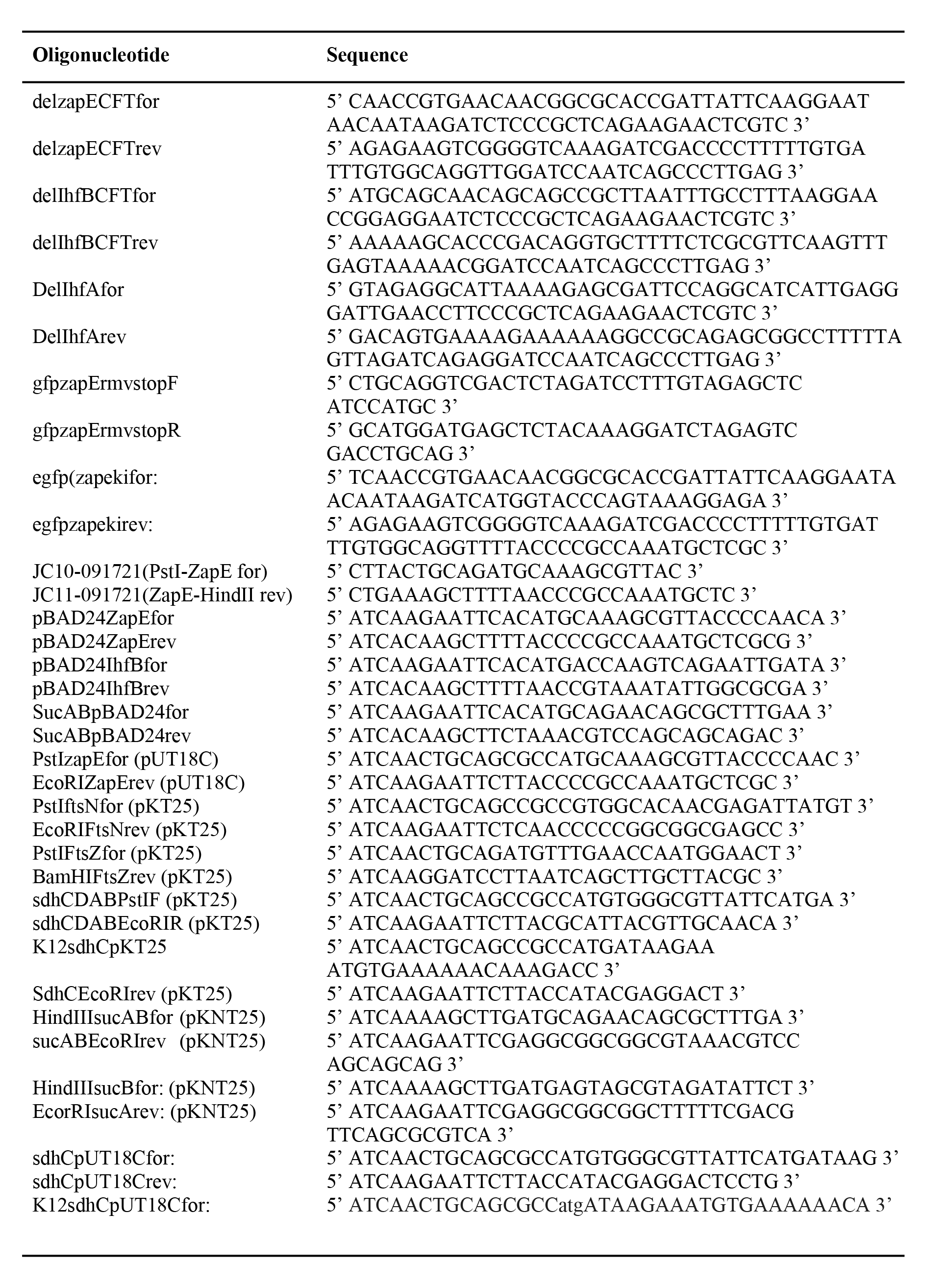
Primer list.

Plasmids pBAD24-*zapE,* pBAD24-*ihfB,* and pBAD24-*sucAB* were constructed by amplifying genes from CFT073 via whole colony PCR using primers pBAD24ZapEfor and pBAD24ZapErev(for pBAD24-*zapE*), pBAD24IhfBfor and pBAD24IhfBrev(for pBAD24-*ihfB*), and SucABpBAD24for and SucABpBAD24rev(for pBAD24-*sucAB*)(table 2). The resulting PCR products were double digested with EcoRI(HiFi) and HindII(HiFi) restriction enzymes and purified via gel purification. Digested PCR products were ligated into EcoRI/HindIII digested pBAD24, transformed into XL10 gold ultracompetent cells (Agilent), and verified by sequencing. Plasmid pBAD24-*gfp_uv3_*-*zapE* was constructed by amplifying *zapE* from CFT073 via whole colony PCR using primers JC10-091721(PstI-ZapE for) and JC11-091721(ZapE-HindII rev) (Table 2). The resulting PCR products were double digested with PstI(HiFi) and EcoRI(HiFi) restriction enzymes and purified via gel purification. Digested PCR products were ligated into PstI/EcoRI digested pBAD24-*gfp_uv3_* (lab stock) (59, 60). Ligation reactions were transformed into XL10 gold ultracompetent cells (Agilent) and verified by sequencing. The stop codon at the end of *gfp* was changed to a glycine by site-directed mutagenesis using primers gfpzapErmvstopF and gfpzapErmvstopR to create pBAD24-*gfp_uv3_*-*zapE*.

Plasmids indicated were then transformed into CFT073, CFT073 Δ*zapE,* or CFT073 Δ*ifhB* strains by electroporation. Strains were grown with ampicillin (50 μg ml^-1^) for the quiescence assay and plated onto 0.2% glucose M9 minimal agar containing ampicillin (50 μg ml^-1^) and L-arabinose (0.1%), where indicated. Lennox broth (LB) supplemented with agar (LB agar) was used for routine cultivation. Liquid and agar 0.2% glucose M9 minimal media were prepared as previously described (29).

Gene deletions were constructed by λred recombination as described by Datsenko and Wanner (42) using primers shown in table 2 and verified by sequencing. Briefly, strains of interest containing pKD46 were grown to an OD_600_ of approximately 0.3 in the presence of ampicillin (100 μg ml^-1^) and L-arabinose (0.36%) at 30 °C. Cells were electroporated with a PCR amplified insert and recombinants were selected for by spread plating onto LB agar supplemented with kanamycin (50 μg ml^-1^) or chloramphenicol (17.5 μg ml^-1^). To construct CFT073 Δ*zapE*::*gfp_uv3_-zapE,* recombinants were selected for by spread plating onto M9 minimal agar containing 1% L-rhamnose. Subsequently, all recombinants were confirmed by sequencing.

### Mini transposon screen and arbitrary PCR

*E. coli* ATM161 (donor) carrying the suicide vector pUT possessing the mini-Tn5 kanamycin (Km) transposon and was conjugated with recipient strain CFT073 (streptomycin resistant) as described previously (29), Both the donor and recipient strain were grown overnight, in LB (Lennox) media. The next day, 100 μl of each culture was mixed in 5 ml of 10 mM MgSO_4_ and filtered through a 0.45 μm nitrocellulose membrane filter, which was subsequently placed on top of a LB (Lennox) agar plate and incubated for 5 h at 37 °C. Filters were vortexed to remove cells in 1 ml glucose M9 minimal media. Cells were centrifuged at 3,000 x *g* for 5 min and washed 2 times with glucose M9 minimal media. Resuspended cells were then diluted, and 10^4^ or 10^5^ CFU ml^-1^ were spread plated onto 0.2% glucose M9 minimal agar with streptomycin (100 μg ml^-1^) and kanamycin (50 μg ml^-1^) and incubated at 37 °C for 30 h. Individual colonies, representing potential CFT073 non-quiescent transposon insertional mutants, were isolated and toothpicked onto glucose M9 minimal agar with streptomycin (100 μg ml^-1^) and kanamycin (50 μg ml^-1^), and were grown overnight at 37 °C. The next day colonies were toothpicked onto LB (Lennox) agar with or without ampicillin (100 μg ml^-1^) and were grown overnight at 37 °C. Colonies that grew on LB (Lennox) agar and demonstrated ampicillin sensitivity, were isolated, cultured, and frozen. Arbitrarily primed PCR for sequencing of transposon insertion was performed precisely as described previously (29).

### Antibiotic tolerance assay

For antibiotic tolerance assays, CFT073, MG1655, or mutant CFT073 overnight cultures, grown in 0.2% glucose M9 minimal medium for 24 h at 37 °C, were diluted to 10^5^ CFU ml^-1^ and 10 ul was spotted onto a 0.22 μm 13 mm filter (Millipore) placed on a M9 minimal with 0.2% glucose agar plate. Plates were incubated for 3 h at 37 °C to induce quiescence, then filters were transferred to an M9 minimal with 0.2% glucose agar plate without antibiotic or with ampicillin (12.5 μg ml^-1^), or nalidixic acid (7.5 μg ml^-1^), and incubated for 20 h at 37 °C. Where indicated, Glucose M9 minimal agar plates were made containing succinate (0.5%) or arabinose (0.1%) and ampicillin (100 μg ml^-1^). Filters were then vortexed in 1 lL of LB to remove cells, serial diluted, spread plate on Luria-Bertani (Lennox) plates, and incubated overnight to obtain viable plate counts. Percent survival was calculated by dividing the number of viable cells on plates containing antibiotic by the number of viable cells on plates without antibiotic. For CFT073 and mutant CFT073 strains tested at high or low cell density, overnight cultures were diluted to an OD_600_ of 0.005 (for low cell density condition) or 0.1 (for high cell density condition) in 10 mls of 0.2% glucose M9 minimal medium. Cultures were then grown at 37 °C for 6 h either shaking (for high cell density condition) or static (for low cell density condition). 10 μl from these cultures was spotted onto a 0.22 μm 13 mm filter (Millipore) on an M9 minimal with 0.2% glucose agar plate with or without ampicillin (12.5 μg ml^-1^) and incubated for 20 h at 37 °C, and then percent survival was calculated as described above.

### Quiescence assay on agar and KGDH assay

Quiescence assay on agar was performed as described previously (29). For KGDH assays, a 24 h overnight culture of CFT073 or mutant strains grown in M9 minimal medium + 0.2% glucose were back diluted into 10 mls of fresh medium to an OD_600_ of 0.01, or an OD_600_ of 0.1 for proliferating CFT073, and grown statically at 37 °C for 20 h. Cultures were then centrifuged to pellet cells, lysed via bacterial protein extraction reagent (B-PER), and protein concentration was determined via addition to Bradford reagent. Protein concentrations were then normalized to 6 μg protein and the KGDH assay kit (Sigma) protocol was followed.

### Bacterial two-hybrid assay (BACTH) X-Gal and Miller assays

The X-Gal based assay was performed by measuring the OD_600_ of overnight cultures grown in LB, and then standardizing the OD_600_ to 1.0 before spotting 10 μls of each culture onto LB kanamycin/ampicillin containing X-Gal (40 μg ml^-1^) and IPTG (0.5 mM) and then incubated at 30 °C. Miller assays were performed by growing cultures overnight in LB, measuring OD_600_, and diluting standardized cultures 1:5 in M9 minimal media + 0.2% glucose. 200 μl of this mixture was added to a 1.2 ml polypropylene 96-well plate and 7 μl of 0.05% sodium dodecyl sulfate (SDS) and 10 μl chloroform was added, and plate was incubated in fume hood shaking for 1 h to evaporate excess chloroform. 100 μl of this mixture was added to 100 μl of a PM2 assay buffer containing 0.1% ortho-Nitrophenyl-β-galactoside (ONPG) and 100 mM β-mercaptoethanol preequilibrated at 37 °C. Reactions were then monitored shaking at 37 °C for yellow color, reactions were stopped by addition of 1 M Na_2_CO_3_ and measured at 415 nm.

### Persister cell assay

24 h overnight cultures were grown in 0.2% glucose M9 minimal medium shaking at 37 °C. by taking a loopful of an LB agar plate streaked with CFT073, CFT073 mutant strain, or MG1655. Cultures were then diluted to an OD_600_ of 0.1 in the presence and absence of ampicillin (100 μg ml^-1^) and grown shaking at 37 °C. Viable plate counts were determined at 0, 2, 4, 6, and 24 h by serial diluting and spread plating onto LB agar plates.

### Microscopy

For visualization of quiescent CFT073, overnight cultures of CFT073 or CFT073 Δ*zapE* were grown at 37 °C for 24 h, back diluted 3 logs, and 5 μl was spotted onto M9 minimal agar + 0.2% glucose. Spots were dried and plates were incubated for 24 h at 37 °C. Cells were resuspended in phosphate-buffered saline (PBS) and applied to an agar gel pad containing 0.2% glucose M9 minimal medium with 5% noble agar.

For visualization of actively growing CFT073, CFT073, mutant strains, or CFT073 strains containing plasmid were struck out on LB agar or LB agar supplemented with ampicillin (100 μg ml^-1^) and grown overnight at 30 °C. A single colony from the plate was then restruck onto LB, or LB supplemented with ampicillin (100 μg ml^-1^) and arabinose (0.01% or 0.2%) and grown for 6 h at 30 °C. Cells were resuspended in phosphate-buffered saline (PBS) and applied to an agar gel pad containing 0.2% glucose M9 minimal medium with 5% noble agar.

Cells were imaged using a Nomarski prism was used to acquire differential interference contrast images, and where indicated, epifluorescence microscopy using an excitation wavelength of 488 nm. Cells were imaged using Zeiss LSM 700 and Nikon Eclipse Ti2 fluorescence microscopes, and images were captured on an AxioCam HRc high-resolution camera with ZEN 2012/NIS-Elements imaging software. Images were processed using Adobe Photoshop CC and analyzed using NIH ImageJ software.

## Acknowledgments

We thank Janet Atoyan for sequencing and microscopy assistance, Ben Piraino, Colby Ferreira, and Cathy Trebino, for helpful suggestions and edits. Microscopy and sequencing were supported by the Rhode Island Institutional Development Award (IDeA) Network of Biomedical Research Excellence from the National Institute of General Medical Sciences of the National Institutes of Health under grant number P20GM103430. We also thank Ben Piraino for assistance in construction of pBAD24-*gfp_uv3_* plasmid.

## Author contributions

J. J. M., E. C. D., D. C. R., P. S. C., and J. L. C. conceptualization; J. J. M., D. A. B., E. K. M., and E. C. D. experimental procedures; J. J. M., D. A. B., E. K. M., and J. L. C. formal analysis; J. J. M. and J. L. C. writing–original draft; J. J. M., and J. L. C. writing–review and editing; E. C. D. resources; J. L. C. visualization; J. L. C. supervision; J. L. C. funding acquisition.

## Funding and additional information

Research reported in this publication was supported in part by the National Institutes of Health under Award Number 1R21AI156574 to J. L. C. The content is solely the responsibility of the authors and does not necessarily represent the official views of the National Institutes of Health or the authors’ respective institution

