## Supporting information for "Metabolic flux regulates growth transitions and antibiotic tolerance in uropathogenic *Escherichia coli*"

**Figure S1. *ihfB* and *zapE* are critical for quiescence and complementation strains respond normally to stimulants.** (A) CFT073 plated at $10^8$ CFU $ml^{-1}$ and CFT073, CFT073 *ihfB* Tn5::kan, CFT073 *zapE* Tn5::kan, CFT073 *gnd* Tn5::kan plated at $10^4$ CFU $ml^{-1}$ on 0.2% glucose minimal agar. (B) CFT073 pBAD24, CFT073 Δ*ihfB* pIhfB, and CFT073 Δ*zapE* pZapE were plated at $10^4$ CFU $ml^{-1}$ on 0.2% glucose minimal agar supplemented with 0.1% arabinose and 50 µg $ml^{-1}$ ampicillin and then challenged with 5 µl of of L-Lys and L-Met (1 mM) or mutanolysin digested *B. subtilis* peptidoglycan (PG). Yellow arrows represent where stimulants were added. Plates in (A) and (B) were incubated at 37 °C for 24 hrs. Pictures in (A) and (B) are representative of at least three independent experiments.

**Figure S2. Antibiotic sensitivity of quiescent and proliferating *E. coli*.** (A) *E. coli* MG1655 and CFT073 inoculated into LB with increasing concentrations of ampicillin measured at 600 nm after 20 h of incubation at 37 °C. (B) CFT073 tolerance to ampicillin (12.5 µg $ml^{-1}$) at low cell density (LCD) and high cell density (HCD) as described in antibiotic tolerance assay in Materials and

Methods. Briefly, strains were diluted to low cell density (0.005 $OD_{600}$) or high cell density (0.1 $OD_{600}$) in 0.2% glucose minimal media and incubated to induce quiescence (for LCD) or growth (for HCD) for 6 h at 37 °C. Cultures were spotted onto 0.22 μm filters on 0.2% glucose minimal agar with and without ampicillin (12.5 μg $ml^{-1}$) and then incubated for 20 h at 37 °C. Viability and percent survival was determined by serial dilution and spread plating. Data in (A) is an average of at least three independent experiments represented as mean ± SEM. Data in (B) is an average of at least four independent experiments represented as mean ± SEM.

**Figure S3. CFT073 Δ*zapE* and ZapE overexpression microscopy and Miller assay confirming X-gal assay results.** (A) DIC microscopy of CFT073 or CFT073 Δ*zapE* strains grown on LB agar and then incubated for 6 h. (B) DIC microscopy of CFT073 or CFT073 Δ*zapE* containing pBAD24 or expressing ZapE from pBAD-ZapE. Cells were incubated for 6 h at 30 °C on LB agar supplemented with ampicillin (100 μg $ml^{-1}$) and arabinose (0.2%). Scale bars in (A) and (B) are 2 μm. (C) BTH101 co-expressing plasmids for X-gal assay plated on LB agar containing ampicillin (100 μg $ml^{-1}$), kanamycin (50 μg $ml^{-1}$), IPTG (0.5 mM), and X-gal (40 μg $ml^{-1}$). Plates were incubated for 24 hrs at 30 °C. (D) Miller assay for β-galactosidase activity performed as described in Materials and Methods. ns signifies not significant, ** signifies a p-value ≤ 0.01, *** signifies a p-value ≤ 0.001, and **** signifies a p-value ≤ 0.0001. Pictures in (C) are representative of at least three independent experiments. Data in (D) is an average of at least five independent experiments represented as mean ± SEM.

**Figure S4. FtsN, ZapE, and SdhC conservation across *E. coli* and ZapE conservation across bacteria.** (A) Clustal Omega alignment of FtsN, ZapE and SdhC amino acid sequences from *E.*

*coli* CFT073, Nissle 1917, or MG1655. (B) Clustal Omega percent identity matrix of ZapE amino acid sequences among various bacterial species.

### Figure S1

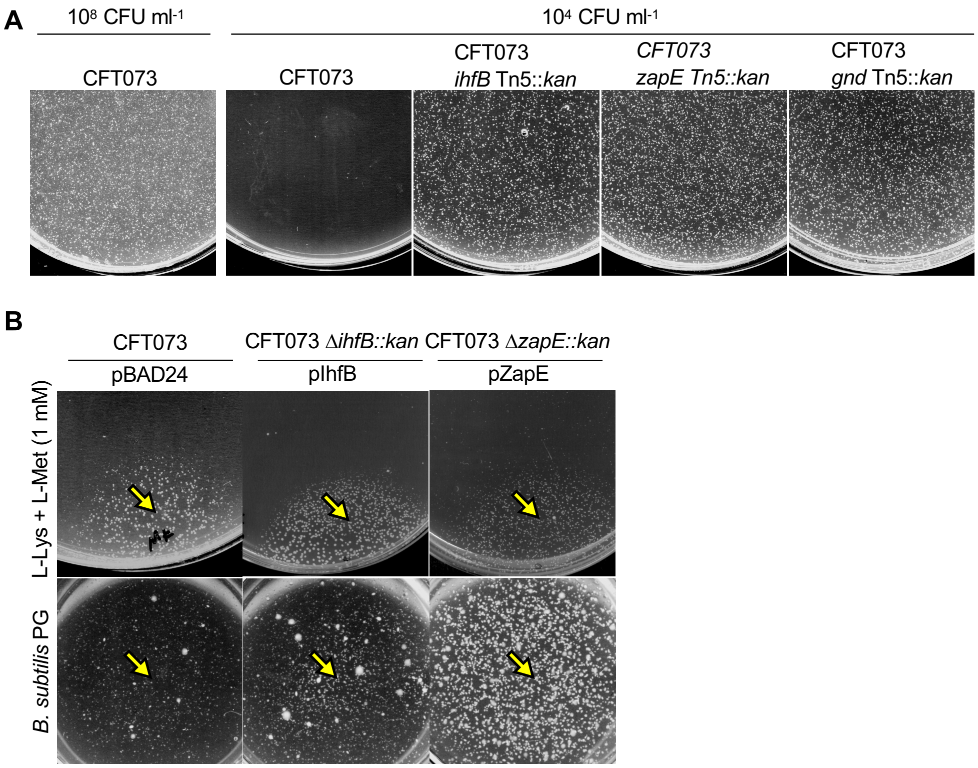
A
$10^8$ CFU $ml^{-1}$
$10^4$ CFU $ml^{-1}$
CFT073
CFT073
CFT073 *ihfB* Tn5::*kan*
*CFT073 zapE Tn5::kan*
CFT073 *gnd* Tn5::*kan*
B
CFT073
CFT073 Δ*ihfB::kan*
CFT073 Δ*zapE::kan*
pBAD24
pIhfB
pZapE
L-Lys + L-Met (1 mM)
*B. subtilis* PG

Figure S2

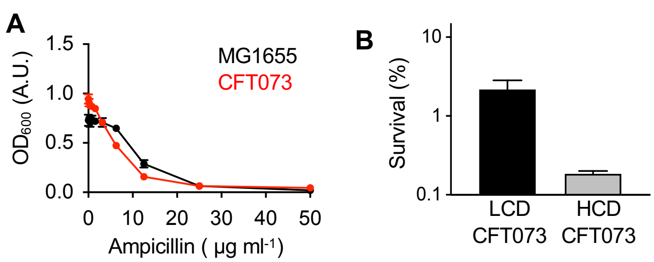
A
1.5
1.0
0.5
0.0
OD600 (A.U.)
MG1655
CFT073
0
10
20
30
40
50
Ampicillin ( µg ml-1)
B
10
1
0.1
Survival (%)
LCD
CFT073
HCD
CFT073

Figure S3

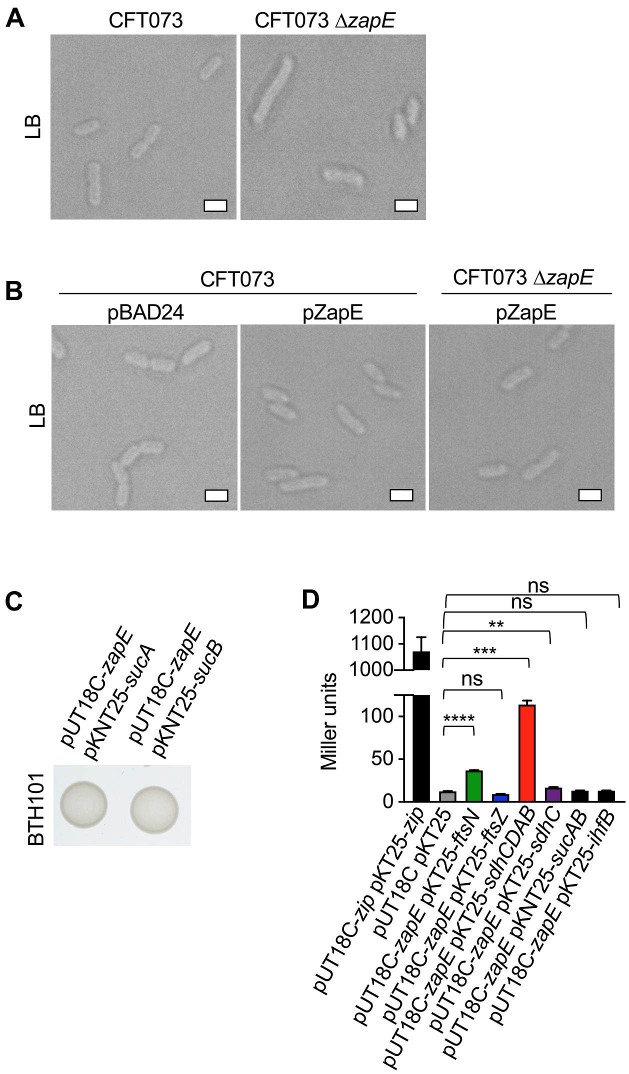
A
CFT073
CFT073 ΔzapE
LB
B
CFT073
CFT073 ΔzapE
pBAD24
pZapE
pZapE
LB
C
pUT18C-zapE pKNT25-sucA
pUT18C-zapE pKNT25-sucB
BTH101
D
Miller units
1200
1100
1000
100
50
0
ns
ns
**
***
ns
****
pUT18C-zip pKT25-zip
pUT18C pKT25
pUT18C-zapE pKT25-ftsN
pUT18C-zapE pKT25-ftsZ
pUT18C-zapE pKT25-sdhCDAB
pUT18C-zapE pKT25-sdhC
pUT18C-zapE pKNT25-sucAB
pUT18C-zapE pKT25-ihfB

### Figure S4

**A**

##### FtsN alignment

```
EcoliCFT073ftsN           MAQRDYVRRSQPAPSRRKKSTSRKKQRNLPAVSPAMVAIAAAVLVTFIGGLYFITHHKKE   60
EcoliNissle1917ftsN       MAQRDYVRRSQPAPSRRKKSTSRKKQRNLPAVSPAMVAIAAAVLVTFIGGLYFITHHKKE   60
EcoliMG1655ftsN           MAQRDYVRRSQPAPSRRKKSTSRKKQRNLPAVSPAMVAIAAAVLVTFIGGLYFITHHKKE   60
                          ************************************************************

EcoliCFT073ftsN           ESETLQSQKVTGNGLPPKPEERWRYIKELESRQPGVRAPTEPSAGGEVKTPEQLTPEQRQ   120
EcoliNissle1917ftsN       ESETLQSQKVTGNGLPPKPEERWRYIKELESRQPGVRAPTEPSAGGEVKTPEQLTPEQRQ   120
EcoliMG1655ftsN           ESETLQSQKVTGNGLPPKPEERWRYIKELESRQPGVRAPTEPSAGGEVKTPEQLTPEQRQ   120
                          ************************************************************

EcoliCFT073ftsN           LLEQMQADMRQQPTQLVEVPWNEQTPEQRQQTLQRQRQAQQLAEQQRLVQQSRTTEQSWQ   180
EcoliNissle1917ftsN       LLEQMQADMRQQPTQLVEVPWNEQTPEQRQQTLQRQRQAQQLAEQQRLVQQSRTTEQSWQ   180
EcoliMG1655ftsN           LLEQMQADMRQQPTQLVEVPWNEQTPEQRQQTLQRQRQAQQLAEQQRLAQQSRTTEQSWQ   180
                          ************************************************.***********

EcoliCFT073ftsN           QQTRTSQAAPVQAQPRQSKPASTQQPYQDLLQTPAHTTAQSKPQQAAPVARVADAPKPTA   240
EcoliNissle1917ftsN       QQTRTSQAAPVQAQPRQSKPASTQQPYQDLLQTPAHTTAQSKPQQAAPVARVADAPKPTA   240
EcoliMG1655ftsN           QQTRTSQAAPVQAQPRQSKPASSQQPYQDLLQTPAHTTAQSKPQQAAPVARAADAPKPTA   240
                          **********************:****************************.********

EcoliCFT073ftsN           EKKDERRWMVQCGSFRGAEQAETVRAQLAFEGFDSKITTNNGWNRVVIGPVKGKENADST   300
EcoliNissle1917ftsN       EKKDERRWMVQCGSFRGAEQAETVRAQLAFEGFDSKITTNNGWNRVVIGPVKGKENADST   300
EcoliMG1655ftsN           EKKDERRWMVQCGSFRGAEQAETVRAQLAFEGFDSKITTNNGWNRVVIGPVKGKENADST   300
                          ************************************************************

EcoliCFT073ftsN           LNRLKMAGHTNCIRLAAGG   319
EcoliNissle1917ftsN       LNRLKMAGHTNCIRLAAGG   319
EcoliMG1655ftsN           LNRLKMAGHTNCIRLAAGG   319
                          *******************
```

##### ZapE alignment

```
EcoliCFT073zapE           MQSVTPTSQYLKALNEGSHQHDDVQKEAVSRLEIIYQELINSTPPAPRTSGLMARVGKLW   60
EcoliNissle1917zapE       MQSVTPTSQYLKALNEGSHQHDDVQKEAVSRLEIIYQELINSTPPAPRTSGLMARVGKLW   60
EcoliMG1655zapE           MQSVTPTSQYLKALNEGSHQPDDVQKEAVSRLEIIYQELINSTPPAPRTSGLMARVGKLW   60
                          ******************** ***************************************

EcoliCFT073zapE           GKREDTKHMPVRGLYMWGGVGRGKTWLMDLFYQSLPGERKQRLHFHRFMLRVHDELTELQ   120
EcoliNissle1917zapE       GKREDTKHMPVRGLYMWGGVGRGKTWLMDLFYQSLPGERKQRLHFHRFMLRVHDELTELQ   120
EcoliMG1655zapE           GKREDTKHTPVRGLYMWGGVGRGKTWLMDLFYQSLPGERKQRLHFHRFMLRVHEELTALQ   120
                          ******** ********************************************:*** **

EcoliCFT073zapE           GQSDPLEIIADRFKAETDVLCFDEFFVSDITDAMLLGGLMKALFARGITLVATSNIPPDE   180
EcoliNissle1917zapE       GQSDPLEIIADRFKAETDVLCFDEFFVSDITDAMLLGGLMKALFARGITLVATSNIPPDE   180
EcoliMG1655zapE           GQTDPLEIIADRFKAETDVLCFDEFFVSDITDAMLLGGLMKALFARGITLVATSNIPPDE   180
                          **:*********************************************************

EcoliCFT073zapE           LYRNGLQRARFLPAIDAIKQHCDVMNVDAGVDYRLRTLTQAHLWLSPLNDETRTQMDKLW   240
EcoliNissle1917zapE       LYRNGLQRARFLPAIDAIKQHCDVMNVDAGVDYRLRTLTQAHLWLSPLNDETRTQMDKLW   240
EcoliMG1655zapE           LYRNGLQRARFLPAIDAIKQHCDVMNVDAGVDYRLRTLTQAHLWLSPLHDETRAQMDKLW   240
                          ************************************************:****:******

EcoliCFT073zapE           LALAGAKRENSPTLEINHRPLATMGVENQTLAVSFTTLCVDARSQHDYIALSRLFHTVML   300
EcoliNissle1917zapE       LALAGAKRENSPTLEINHRPLATMGVENQTLAVSFTTLCVDARSQHDYIALSRLFHTVML   300
EcoliMG1655zapE           LALAGGKRENSPTLEINHRPLATMGVENQTLAVSFTTLCVDARSQHDYIALSRLFHTVML   300
                          *****.******************************************************

EcoliCFT073zapE           FDVPVMTRLMESEARRFIALVDEFYERHVKLVVSAEVPLYEIYQGERLKFEFQRCLSRLQ   360
EcoliNissle1917zapE       FDVPVMTRLMESEARRFIALVDEFYERHVKLVVSAEVPLYEIYQGERLKFEFQRCLSRLQ   360
EcoliMG1655zapE           FDVPVMTRLMESEARRFIALVDEFYERHVKLVVSAEVPLYEIYQGDRLKFEFQRCLSRLQ   360
                          **********************************************:*************

EcoliCFT073zapE           EMQSEEYLKREHLAG   375
EcoliNissle1917zapE       EMQSEEYLKREHLAG   375
EcoliMG1655zapE           EMQSEEYLKREHLAG   375
                          ***************
```

##### SdhC alignment

```
EcoliCFT073sdhC           MWALFMIRNVKKQRPVNLDLQTIRFPVTAIASILHRVSGVITFVAVGILLWLLGTSLSSP   60
EcoliNissle1917sdhC       MWALFMIRNVKKQRPVNLDLQTIRFPVTAIASILHRVSGVITFVAVGILLWLLGTSLSSP   60
EcoliMG1655sdhC           -----MIRNVKKQRPVNLDLQTIRFPITAIASILHRVSGVITFVAVGILLWLLGTSLSSP   55
                               **********************:********************************

EcoliCFT073sdhC           EGFEQASAIMGSFFVKFIMWGILTALAYHVVVGIRHMMMDFGYLEETFEAGKRSAKISFV   120
EcoliNissle1917sdhC       EGFEQASAIMGSFFVKFIMWGILTALAYHVVVGIRHMMMDFGYLEETFEAGKRSAKISFV   120
EcoliMG1655sdhC           EGFEQASAIMGSFFVKFIMWGILTALAYHVVVGIRHMMMDFGYLEETFEAGKRSAKISFV   115
                          ************************************************************

EcoliCFT073sdhC           ITVVLSLLAGVLVW   134
EcoliNissle1917sdhC       ITVVLSLLAGVLVW   134
EcoliMG1655sdhC           ITVVLSLLAGVLVW   129
                          **************
```

**B**

| Percent AA sequence identity | *E. coli CFT073* | *E. coli MG1655* | *S. Flexneri* | *S. enterica* | *V. cholerae* | *B. cenocipacia* | *P. aeruginosa* | *N. gonorrhoeae* | *K. pneumoniae* |
|---|---|---|---|---|---|---|---|---|---|
| *E. coli CFT073* | 100 | 97.6 | 97.6 | 85.03 | 53.83 | 44.01 | 39.66 | 38.84 | 30.49 |
| *E. coli MG1655* | 97.6 | 100 | 98.67 | 85.03 | 53.28 | 43.73 | 39.94 | 38.84 | 30.79 |
